# Whole-genome sequencing data of a diverse grapevine germplasm collection maintained in Bordeaux, France

**DOI:** 10.64898/2026.07.31.742002

**Authors:** Marina de Miguel, Maria Lafargue, Enrique Sáez-Laguna, Joseph Tran, Nabil Girollet, Pierre-François Bert, Yi Wang, Zhenchang Liang, Sabine Guillaumie, Zhanwu Dai, Nathalie Ollat

## Abstract

Grapevine (*Vitis vinifera*) is one of the world’s most economically important fruit crops and a model species for perennial fruit tree genetics and genomics. The extensive genetic diversity found in cultivated and wild *Vitis* species provides a valuable resource for studies of domestication, adaptation, trait evolution, and breeding. This article presents a standardized whole-genome variant dataset comprising 547 grapevine accessions maintained in the INRAE Bordeaux grapevine germplasm collection, including 397 domesticated *V. vinifera* cultivars and 150 wild *Vitis* accessions. Whole-genome sequencing data were generated at a target sequencing depth of approximately 20×, and sequence variants were identified using a standardized Genome Analysis Toolkit (GATK) workflow against the reference genome PN40024v4 (40X). Variant discovery was carried out simultaneously across the complete sample set to ensure consistent genotype calling.

All accessions were sequenced using the same technology and processed using the same reference genome, sequence alignment, variant-calling, and filtering workflow to produce a standardized variant dataset comprising ca. 9.1M SNPs and 0.77M INDELs. The resulting VCF files provide a harmonized genomic resource that can be readily reused for studies of grapevine genetics, germplasm characterization, population genomics, comparative genomics, genome-wide association studies, and the development and benchmarking of bioinformatic methods.

**SPECIFICATIONS TABLE:** 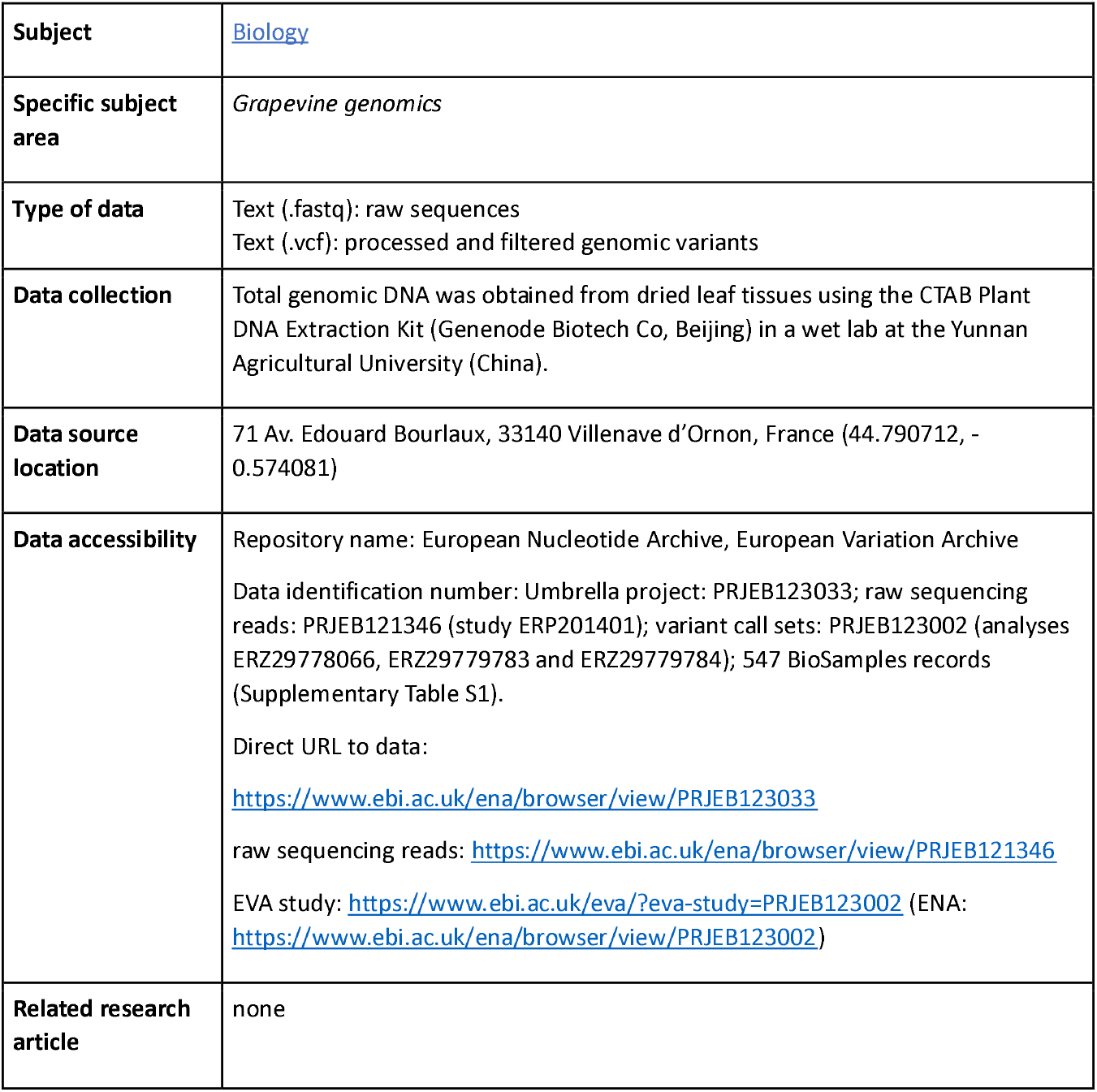

**VALUE OF THE DATA:**

- This dataset provides whole-genome raw sequencing for 118 wild *Vitis* accessions originating from North America and Asia and a sequencing-derived variant data for these accessions and 429 *Vitis vinifera* cultivars and wild accessions from the INRAE Bordeaux germplasm collection, previously published by Dong et al. 2023[1]. The dataset captures genetic variation across a total of 547 grapevine accessions, including domesticated and wild grapevine germplasm using a common variant-calling pipeline, facilitating direct comparisons among accessions.
- The inclusion of wild *Vitis* species together with cultivated grapevine accessions provides a resource for studies of grapevine diversity, domestication, phylogenetic relationships, and comparative genomics. The dataset enables the investigation of genetic variation across multiple *Vitis* species using a standardized set of genomic variants.
- The variant call format (VCF) file can be reused for population genetics, phylogenetic analyses, genetic diversity assessments, introgression analyses, and the identification of genomic regions of interest. The dataset is compatible with widely used bioinformatics software and can be integrated with other publicly available grapevine genomic resources.
- This dataset constitutes a genomic resource for grapevine breeding and conservation research. Researchers can use these data to identify genetic diversity in wild relatives, compare allelic variation between cultivated and wild germplasm, investigate candidate loci associated with traits of interest, and support the management and characterization of grapevine germplasm collections.

## BACKGROUND

The grapevine germplasm collection maintained by INRAE at Bordeaux consists of cultivars originated and cultivated in different regions throughout the world and their wild relatives. Whole genome sequencing of the accessions belonging to this collection was conceived to support genomic studies of grapevine diversity by providing whole-genome variant data for domesticated *Vitis vinifera* cultivars and wild *Vitis* species. The original motivation for generating part of these data is described by Dong et al. (2023)[1], who investigated grapevine domestication and the origin of domestication-related traits using whole-genome sequencing of cultivated and *Vitis vinifera ssp. sylvestris* accessions from Eurasia. The present dataset expands on the sequencing data reported by Dong et al. (2023) [1]], which included cultivated grapevine (*Vitis vinifera*) and *V. vinifera* ssp. *sylvestris* accessions from the INRAE Bordeaux germplasm collection. In addition to these accessions, the present release incorporates whole-genome sequencing data from 118 wild *Vitis* accessions representing species native to North America and Asia from the same collection. Unlike the previous study, which released the raw sequencing data and generated variants relative to a *V. vinifera* ssp. *sylvestris* reference genome, the present dataset provides a unified, ready-to-use variant call format (VCF) file generated through joint variant calling against the widely used PN40024 reference genome [2]. This standardized variant dataset facilitates integration with existing grapevine genomic resources and enables comparative analyses across cultivated and wild *Vitis* species without requiring users to perform computationally intensive variant calling.

The inclusion of genomic variants from wild North American and Asian *Vitis* accessions combined with the previously available *V. vinifera ssp*. sativa and ssp. sylvestris represents a valuable resource for breeding. In this sense, North American Vitis spp. are particularly important for breeding resistant varieties to the main grapevine pathogens (i.e. powdery mildew and downy mildew)[3] and for rootstock breeding because most of them are resistant to phylloxera[4]. Since the widespread adoption of grafting onto resistant rootstocks following the phylloxera crisis of the late nineteenth century, rootstock breeding has relied on a remarkably narrow genetic base, primarily involving a few North American *Vitis* species (*V. berlandieri, V. riparia, V. rupestris, V. champini*) and, to a lesser extent *V. vinifera*. However, the genus *Vitis* comprises approximately 70 species distributed across North America, Europe, and Asia, most of which are interfertile and remain largely unexplored for rootstock improvement[5].

## DATA DESCRIPTION

This dataset comprises whole-genome variants: Single Nucleotide Polymorphims (SNPs) and INDELs for 547 grapevine accessions, including 397 *Vitis vinifera* cultivars and 150 wild *Vitis* accessions. Grapevine wild relatives in this dataset belong to 18 species native to North America, six to Asia and the Eurasian species *Vitis vinifera ssp. sylvestris* (Table 1). These variants are provided as six compressed Variant Call Format (VCF) files, one per variant type (SNPs, INDELs) and genomic compartment (nuclear, chloroplast and mitochondrion).

**Table 1.**
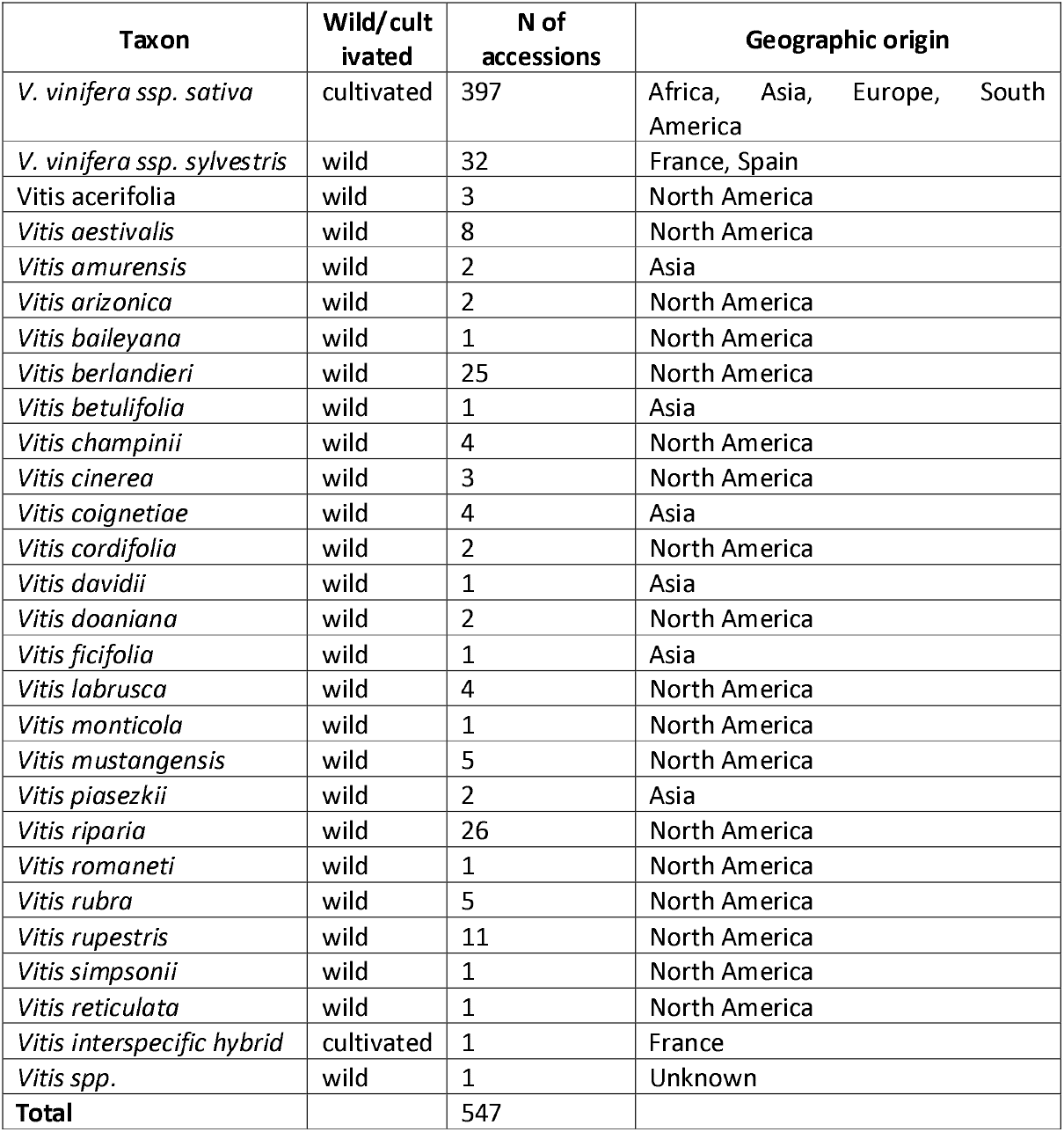
Summary of the samples in the dataset.

| Taxon | Wild/cultivated | N of accessions | Geographic origin |
| --- | --- | --- | --- |
| *V. vinifera ssp. sativa* | cultivated | 397 | Africa, Asia, Europe, South America |
| *V. vinifera ssp. sylvestris* | wild | 32 | France, Spain |
| *Vitis acerifolia* | wild | 3 | North America |
| *Vitis aestivalis* | wild | 8 | North America |
| *Vitis amurensis* | wild | 2 | Asia |
| *Vitis arizonica* | wild | 2 | North America |
| *Vitis baileyana* | wild | 1 | North America |
| *Vitis berlandieri* | wild | 25 | North America |
| *Vitis betulifolia* | wild | 1 | Asia |
| *Vitis champinii* | wild | 4 | North America |
| *Vitis cinerea* | wild | 3 | North America |
| *Vitis coignetiae* | wild | 4 | Asia |
| *Vitis cordifolia* | wild | 2 | North America |
| *Vitis davidii* | wild | 1 | Asia |
| *Vitis doaniana* | wild | 2 | North America |
| *Vitis ficifolia* | wild | 1 | Asia |
| *Vitis labrusca* | wild | 4 | North America |
| *Vitis monticola* | wild | 1 | North America |
| *Vitis mustangensis* | wild | 5 | North America |
| *Vitis piasezkii* | wild | 2 | Asia |
| *Vitis riparia* | wild | 26 | North America |
| *Vitis romaneti* | wild | 1 | North America |
| *Vitis rubra* | wild | 5 | North America |
| *Vitis rupestris* | wild | 11 | North America |
| *Vitis simpsonii* | wild | 1 | North America |
| *Vitis reticulata* | wild | 1 | North America |
| *Vitis interspecific hybrid* | cultivated | 1 | France |
| *Vitis spp.* | wild | 1 | Unknown |
| **Total** |  | 547 |  |

The filtered dataset contains approximately 9.1 million SNPs and 0.77M INDELs (Table 2). They are distributed across the 19 grapevine chromosomes, the chloroplastic and mitochondrial genomes (Figure 1). All the variants are reported relative to the PN40024v4 (40X)[2] grapevine reference genome. The VCF files follow the standard VCF specification and include chromosome identifier, genomic position, reference and alternate alleles, variant quality metrics, filtering status, INFO annotations, FORMAT descriptors, and genotype information for each accession.

**Table 2.** Summary statistics of the genomic variants dataset.

| Metric | SNPs | INDELs |
| --- | --- | --- |
| Number of samples | 547 | 547 |
| Wild samples | 150 | 150 |
| Cultivated samples | 397 | 397 |
| Species represented | 25 | 25 |
| Number of variants | 9 092 040 | 769 347 |
| Mean variants per chromosome | 478 528 | 40 492 |
| Number of variants in the chloroplastic genome | 74 | 26 |
| Number of variants in the mitochondrial genome | 498 | 244 |
| Mean missing genotype rate | 0.0335 | 0.0295 |
| Mean minor allele frequency | 0.1801 | 0.1634 |
| Mean genotype quality | 40.26 | 40.58 |
| Mean read depth at variant sites | 12.2X | 11.8X |

**Figure 1.**
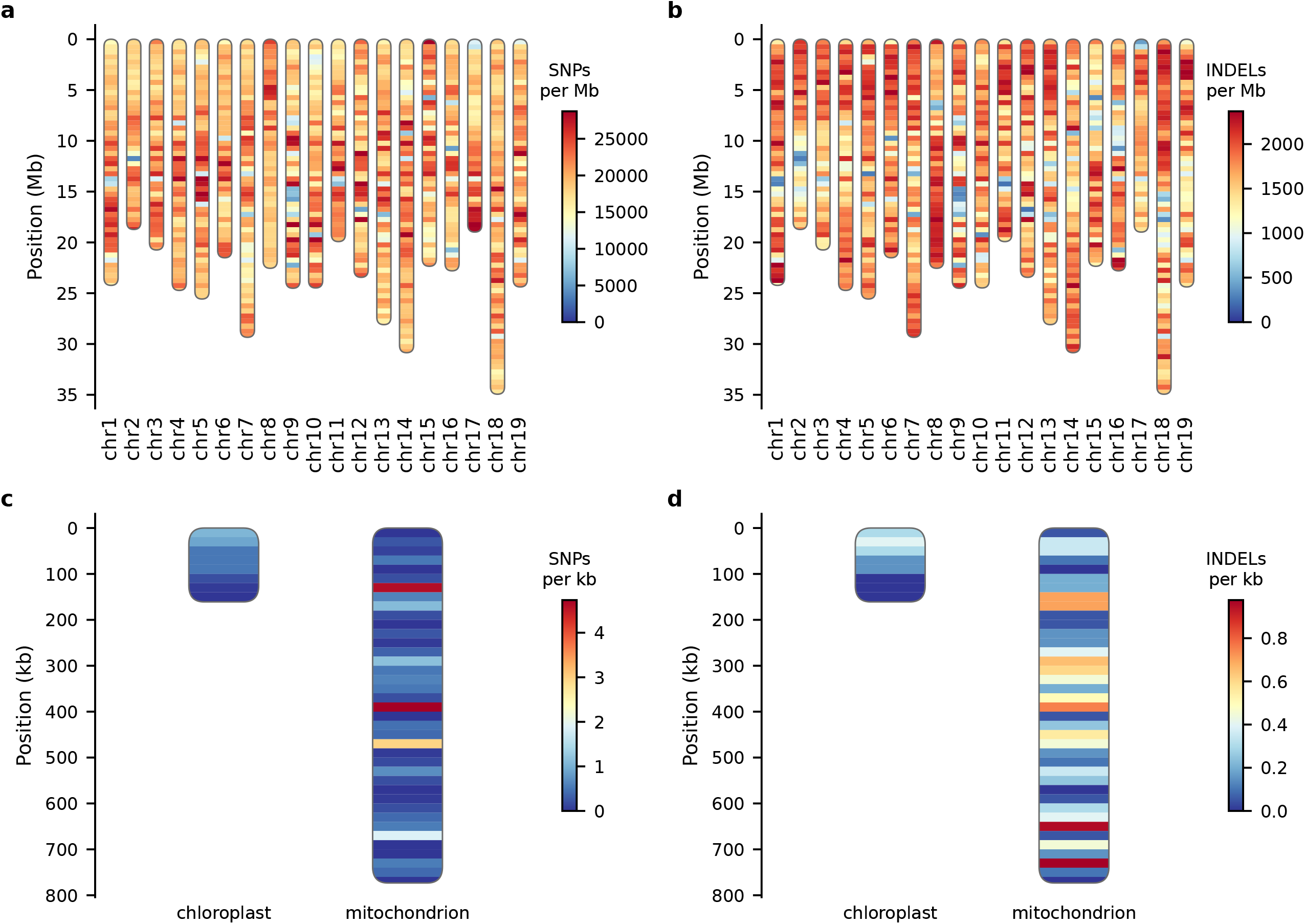
Variant density ideograms per chromosome and compartment: a) mean SNP density per 500kb window in the nuclear genome; b) mean INDEL density per 500kb window in the nuclear genome; c) mean SNP density per 20kb window in the organelle genomes; d) mean INDEL density per 20kb window in the organelle genomes. Centromeres could not be positioned because the information was not available in the PN40024v4 (40X) reference genome.

The complete list of accessions included in the variant calling of this study, as well as the Vitis Information Variety Catalog (VIVC) information, is presented in Supplementary Table S1.

All data are archived at EMBL-EBI and can be accessed through a single umbrella project, PRJEB123033, which groups two child projects: PRJEB121346 for the raw sequencing reads and PRJEB123002 for the variant call sets. Both refer to the same 547 BioSamples records.

### Raw sequencing reads

Paired-end whole-genome sequencing reads for the 118 newly sequenced wild *Vitis* accessions are available in the European Nucleotide Archive under project PRJEB121346 (study ERP201401); individual run accessions are listed in Supplementary Table S1. Reads for the 429 previously published accessions are not redistributed here; they remain available from the original study of Dong *et al*. in the Genome Sequence Archive under accession CRA006917 (BioProject PRJCA009314).

### Variant call sets

Filtered variants for the 547 accessions are available in the European Variation Archive under project PRJEB123002, as gzip-compressed VCF files, one per variant type and genomic compartment: eva_SNP_nuclear.vcf.gz and eva_INDEL_nuclear.vcf.gz for the 19 nuclear chromosomes (reference GCA_927798565, PN40024.v4; analysis accession ERZ29778066), eva_SNP_chloroplast.vcf.gz and eva_INDEL_chloroplast.vcf.gz for the chloroplast genome (reference DQ424856.1; analysis accession ERZ29779783), and eva_SNP_mitochondrion.vcf.gz and eva_INDEL_mitochondrion.vcf.gz for the mitochondrial genome (reference FM179380.1; analysis accession ERZ29779784). Each VCF file is accompanied by its tabix index (.tbi), and sample columns are named consistently across all files. Sample columns follow the naming scheme <collection number>-<species>-<accession name>, which is the identifier used to link each genotype column to the corresponding entry in Supplementary Table S1.

### Sample records

The 547 accessions are described by 547 BioSamples records, each cross-referenced either to the corresponding ENA run (newly sequenced accessions) or to the original GSA sample (reused accessions). The complete correspondence between accession name, species, BioSample accession and sequencing data is provided in Supplementary Table S1. Supplementary Table S1 is therefore the key that links the three components of the dataset: for each of the 547 accessions it gives the sample name used in the VCF columns, the species and accession name, the BioSample accession and, for the 118 newly sequenced accessions, the corresponding ENA experiment and run accessions.

## EXPERIMENTAL DESIGN, MATERIALS AND METHODS

Whole-genome sequencing data were generated as described in Dong et al. 2023 [1]. Briefly, total genomic DNA obtained from dried grapevine leaf tissues using the CTAB Plant DNA Extraction Kit (Genenode Biotech Co, Beijing) in a wet lab at the Yunnan Agricultural University (China). Sequencing libraries with an insert size of 350~550 bp were prepared with NEBNext® UltraTM DNA Library Prep Kit (Illumina, USA) according to the manufacturer’s directions. Paired-end sequencing was performed on an Illumina NovaSeq 6000 platform by both Novogene (Beijing, China) and Berry Genomics (Beijing, China). The target sequencing depth was 20x for each accession.

The obtained sequences were aligned against the reference genome PN40024v4 (40X)[2]. Individual genomic variant call (gVCF) files produced with the Genome Analysis Toolkit (GATK) HaplotypeCaller [6] were used as the starting point for this work (see Figure 1 for the analysis pipeline).

Joint variant calling was performed following the GATK Best Practices workflow. Individual gVCF files were combined using *GenomicsDBImport*, followed by joint genotyping using *GenotypeGVCFs* against the reference genome. Variant discovery was carried out simultaneously across the complete sample set to ensure consistent genotype calling.

Variant sites were extracted from invariant positions using GATK SelectVariants. SNPs and INDELs were then filtered independently using the GATK hard-filtering recommendations. SNP filtering included Quality by Depth (QD <2.0), variant quality (QUAL <30.0), Fisher Strand (FS >60.0), Strand Odds Ratio (SOR >3.0), Mapping Quality (MQ <40.0), MQRankSum <-12.5, and ReadPosRankSu <-8.0 metrics. INDELs were filtered using QD, QUAL, FS, and ReadPosRankSum metrics. Variants with more than 20% missing genotypes across accessions were removed from the final dataset. In addition, the SNPs dataset was filtered for biallelic SNPs with Minor Allele Frequency (MAF) higher or equal to 0.05. For the dataset deposited in the European Variation Archive, the same MAF ≥ 0.05 threshold was subsequently applied to the INDEL dataset. The variant counts reported in Tables 2 and 3 therefore refer to this deposited dataset.

The filtered VCF files were compressed using *BGZF* and indexed with *Tabix* to facilitate downstream analyses.

All analyses were performed on the Genotoul High-Performance Computing platform (Toulouse, France).

Supplementary quality-control analyses were done to characterize the genomic distribution of missing genotypes and their relationship with sequence mappability and repeat content. Genome mappability (GenMap v1.3.0, k = 150 nt, ≤ 2 mismatches) and repeat content (RepeatMasker v4.1.5 / RMBlast with a plant RepBase library 27.02 limited to Viridiplantae/Eudicot) were computed per 500-kb window on PN40024.v4[2]. The per-window excess of missing genotypes in wild non-vinifera accessions was correlated (Spearman) with both, and compared between the most- and least-affected windows (see supplementary Figure S1).

## LIMITATIONS

The dataset has several limitations that should be considered when reused. Although it includes a broad representation of cultivated *Vitis vinifera* and wild *Vitis* species, the number of accessions available for most wild species is limited, which restricts analyses of intraspecific genetic diversity and population structure. In addition, the sampled accessions do not cover the complete geographic distribution of each species, and therefore may not capture their full genetic diversity or regional population differentiation.

Across the newly sequenced accessions, 98.8 % of reads mapped to the PN40024.v4 reference with a mean PCR duplication rate of 13.3 %, giving a mean mapped coverage of 17.3× (range 12.6–24.8×), or 15.0× after duplicate removal. Comparable values were obtained for the reused accessions, with mean coverage after duplicate removal of 15.4× for *V. vinifera ssp. sylvestris*. While this depth is suitable for reliable SNP discovery and genotyping, it may limit applications requiring higher sequencing depth, such as the detection of low-frequency variants, somatic mutations, complex structural variants, or analyses requiring highly accurate genotype calls in repetitive genomic regions. In addition, the mean per-sample depth of the complete variant dataset (n=547 accessions) after read filtering is 12.2× for SNPs and 11.8× for INDELs, indicating that the depth achieved is below the initial target.

The proportion of variant sites with no read coverage is markedly higher for wild Vitis accessions belonging to species phylogenetically distant from the *V. vinifera* reference genome (Figure 3). Consequently, missing genotypes are structured by species and concentrated in specific genomic regions, which should be taken into account in comparative analyses across the panel. Supplementary analyses further show that these regions are characterized by low sequence mappability and an enrichment of repetitive elements, particularly LTR retrotransposons (Supplementary Figure S1), consistent with reduced read alignment to the *V. vinifera* reference genome. In addition, the excess of missing genotypes in wild accessions is highly correlated between SNPs and INDELs genome-wide (Supplementary Figure S1), indicating that this pattern is consistent across variant types.

## ETHICS STATEMENT

The authors have read and follow the ethical requirements for publication in Data in Brief. The current work does not involve human subjects, animal experiments, or any data collected from social media platforms.

## CREDIT AUTHOR STATEMENT

**Marina de Miguel**: Supervision, Writing-Original Draft. **Maria Lafargue**: Resources, Data Curation. **Enrique Sáez-Laguna**: Investigation, Formal Analysis, Methodology. **Joseph Tran**: Visualization, Writing-Review & Editing. **Nabil Girollet**: Formal Analysis. **Pierre-François Bert**: Methodology. **Yi Wang**: Investigation, Writing-Review & Editing. **Zhenchang Liang**: Investigation, Writing-Review & Editing. **Sabine Guillaumie**: Data curation, Writing-Review & Editing. **Zhanwu Dai**: Investigation, Conceptualization, Funding Acquisition, Writing-Review & Editing. **Nathalie Ollat**: Conceptualization, Funding Acquisition, Writing-Review & Editing.

## ACKNOWLEDGEMENTS

This work was supported by Grape Sequencing and Re-sequencing project, LIA INNOGRAPE II International Associated Laboratory, Chinese Academy of Sciences President’s International Fellowship Initiative (Grant No. 2025PG0014), PHC CAI YUANPEI No. 52510VJ.

We gratefully acknowledge Louis Bordenave for his contributions to the maintenance, curation, and study of the germplasm collection. We also thank Nicolas Hocquard, Cyril Hevin, Jean-Pierre Petit, and Aurélien Deloume for maintaining the experimental vineyards. The authors acknowledge the INRAE Experimental Unit “Vigne et Vin Bordeaux Grande Ferrade” (UE 1442) for maintaining the experimental vineyards and providing technical support. We gratefully acknowledge the INRAE CRB Vigne Vassal for their support in maintaining the Bordeaux germplasm collection.

We are grateful to the genotoul bioinformatics platform Toulouse Occitanie (Bioinfo Genotoul, https://doi.org/10.15454/1.5572369328961167E12) for providing help, computing and storage resources.

## SUPPLEMENTARY MATERIAL

**Table S1. List of grapevine accessions sequenced in the variant calling of this study**. VIVC database last accessed on July 17th of 2026. According to VIVC periodic updates, some information presented in this table may differ in future requests.

**Supplementary Figure S1**. Genomic features associated with regions enriched in missing genotypes in wild Vitis accessions. (a) Relationship between genome mappability (GenMap, k = 150) and repeat fraction per 500-kb window; points are coloured by the excess of missing genotypes in wild Vitis spp. relative to V. vinifera accessions (Spearman rho = −0.64). (b) Correlation between the excess of missing genotypes computed independently on SNPs and on INDELs across 500-kb windows (Spearman rho = 0.90, R2 = 0.88, n = 933); windows in the top 5% of SNP excess are highlighted. (c) Mean fraction of each 500-kb window occupied by the different classes of repetitive elements, in windows with the highest (HOT, top 5%) versus lowest (COLD, bottom 5%) excess of missing genotypes. (d) Enrichment of each repeat class in HOT versus COLD windows (Mann-Whitney test; *** p < 0.001, ns not significant); the Satellite class is omitted as it occupies less than 0.1% of the genome. Excess values are expressed in percentage points (pp).

## Declaration of generative AI and AI-assisted technologies in the manuscript preparation process statement

During the preparation of this work, the authors used ChatGPT to create Figure 2 and Claude to assist with to assist with script writing. After using these tools, the authors reviewed and edited the content as needed and take full responsibility for the content of the published article.

**Figure 2.**
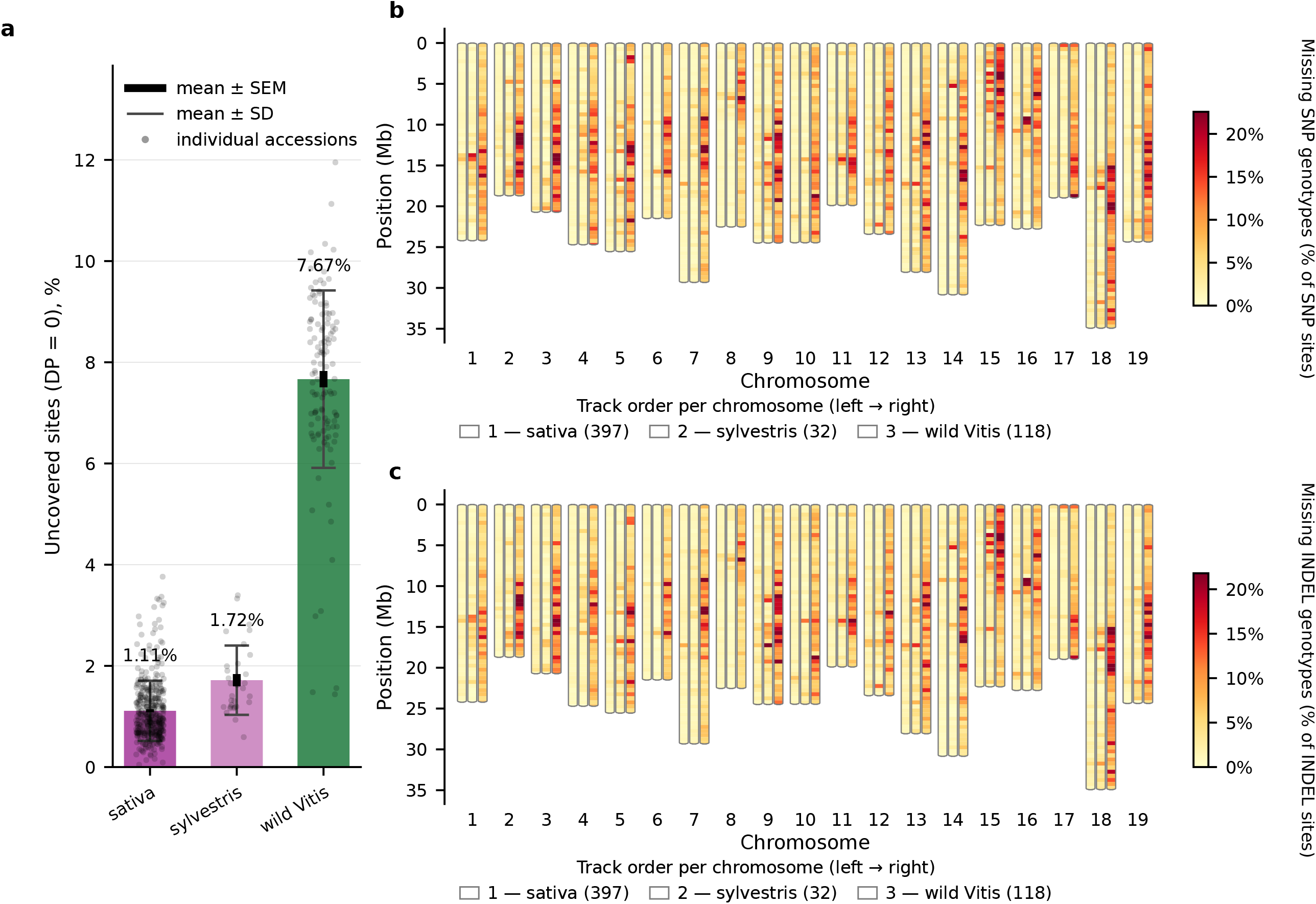
Workflow used to generate the variant dataset, from whole-genome sequencing to the production of the filtered VCF files.

**Figure 3.**
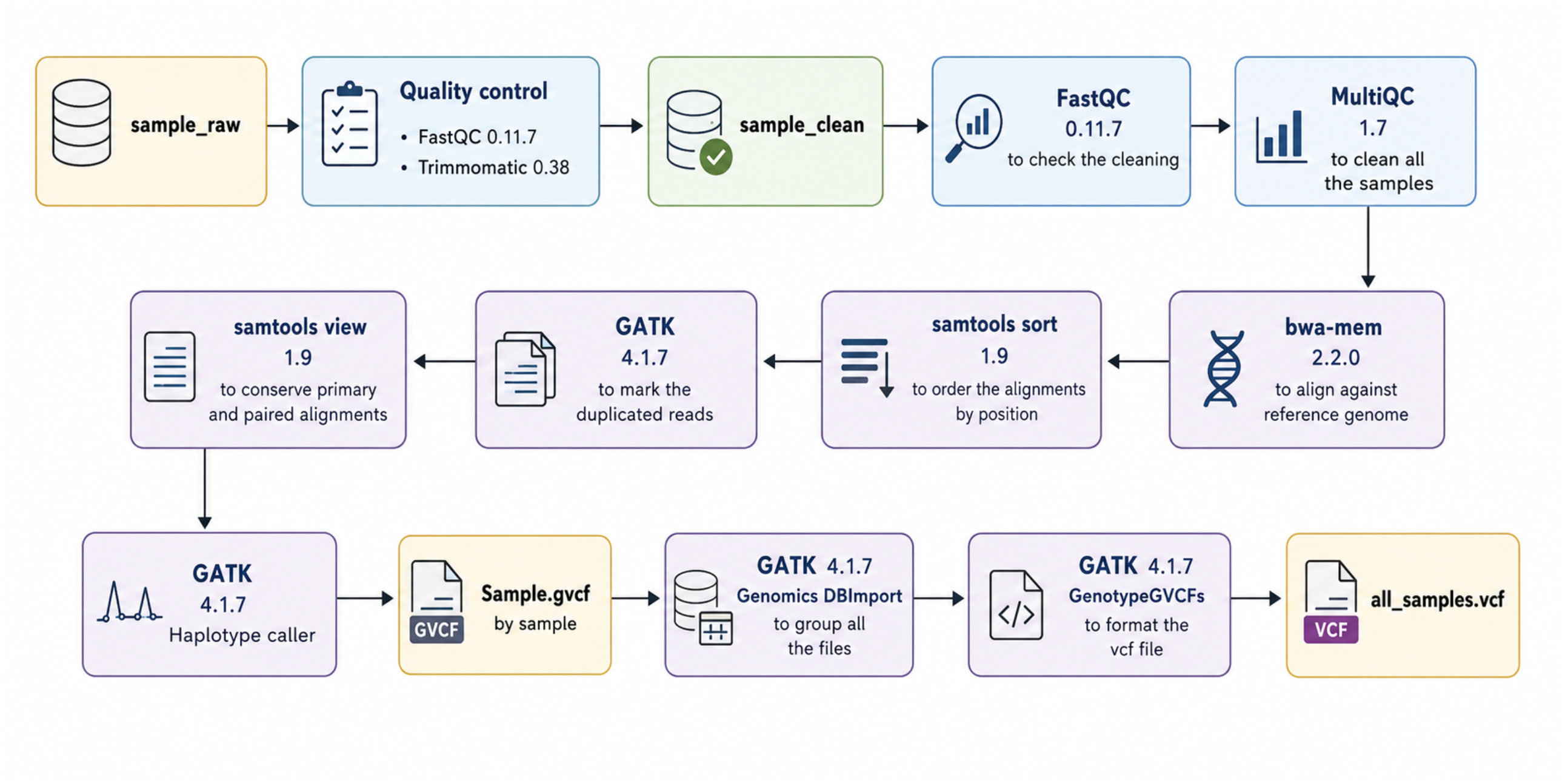
Uncovered variants distribution; a) per group of species: cultivated, wild *V. vinifera spp. sylvestris*, other wild *Vitis spp*. b) mean missing SNP genotypes per 500kb window in the nuclear genome; c) mean missing INDEL genotypes per 500kb window in the nuclear genome.

