## Supplementary figures and images for "Whole-genome sequencing data of a diverse grapevine germplasm collection maintained in Bordeaux, France"

### Supplmentary Figure S1

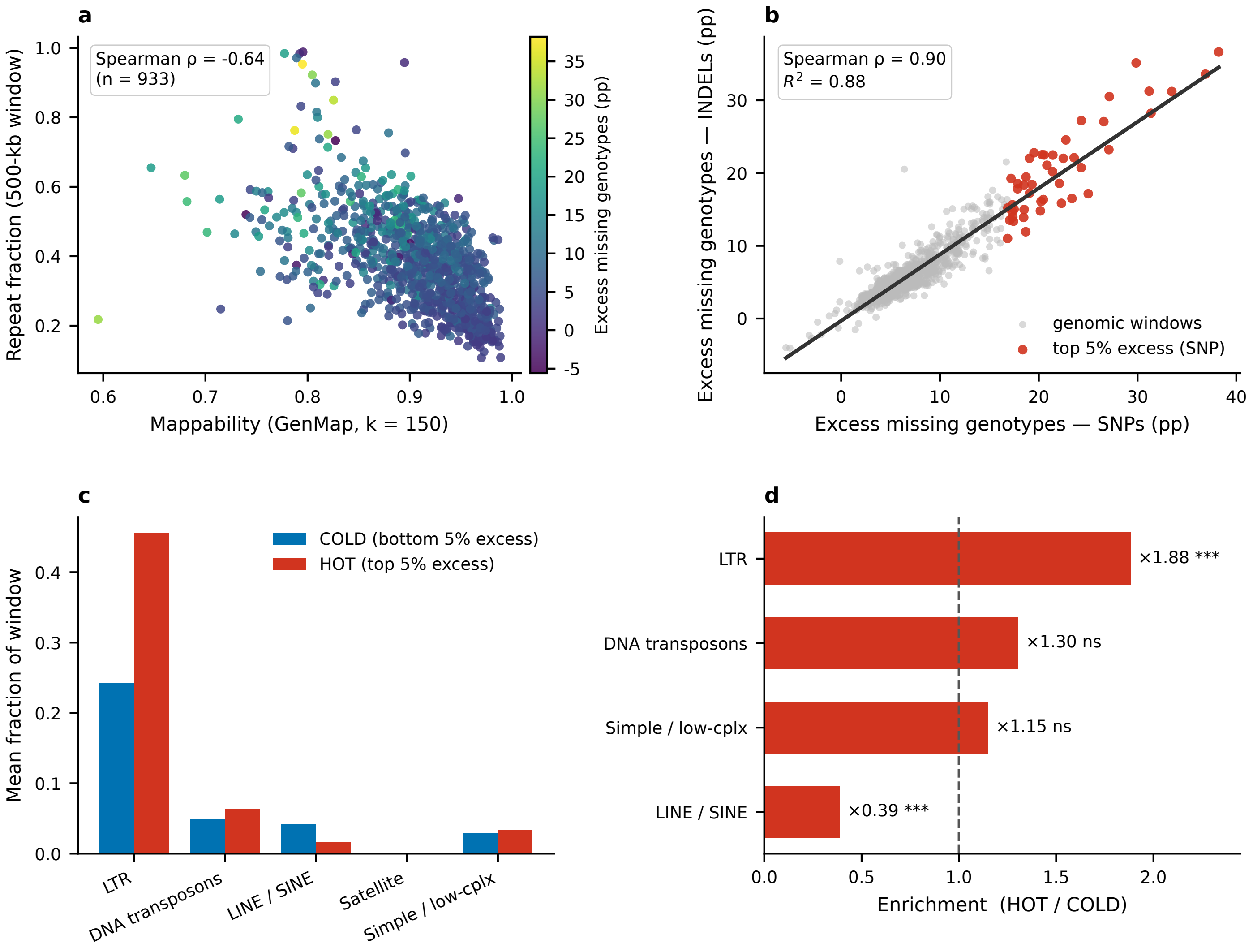
